# Short DNA sequence patterns accurately identify broadly active human enhancers

**DOI:** 10.1101/111955

**Authors:** Laura L. Colbran, Ling Chen, John A. Capra

## Abstract

Enhancers are DNA regulatory elements that influence gene expression. There is substantial diversity in enhancers’ activity patterns: some enhancers drive expression in a single cellular context, while others are active across many. Sequence characteristics, such as transcription factor (TF) binding motifs, influence the activity patterns of regulatory sequences; however, the regulatory logic through which specific sequences drive enhancer activity patterns is poorly understood. Recent analysis of *Drosophila* enhancers suggested that short dinucleotide repeat motifs (DRMs) are general enhancer sequence features that drive broad regulatory activity. However, it is not known whether the regulatory role of DRMs is conserved across species. We performed a comprehensive analysis of the relationship between short DNA sequence patterns, including DRMs, and human enhancer activity in 38,538 enhancers across 411 different contexts. In a machine-learning framework, the occurrence patterns of short sequence motifs accurately predicted broadly active human enhancers. However, DRMs alone were weakly predictive of broad enhancer activity in humans and showed different enrichment patterns than in *Drosophila*. In general, GC-rich sequence motifs were significantly associated with broad enhancer activity; consistent with this enrichment, broadly active human TFs recognize GC-rich motifs. Our results reveal the importance of specific sequence motifs in broadly active human enhancers, demonstrate the lack of evolutionary conservation of the role of DRMs, and provide a computational framework for investigating the logic of enhancer sequences.

## INTRODUCTION

Enhancers are DNA regulatory elements distal to promoters that bind transcription factors (TFs) to drive tissue-specific gene expression. They control patterns of gene expression during development, allowing diverse tissues to differentiate from a single cell and continue functioning properly in maturity (1,2). Because enhancers play a central role in regulating essential transcriptional programs, genome-wide association studies (GWAS) often implicate non-coding variation in enhancer regions as associated with risk for numerous complex diseases (3,4). Several in-depth experimental analyses of loci identified by GWAS have revealed that the causal mutations in these regions disrupt enhancer activity (5–8). However, the function of many of these variants is unknown, and it can be unclear in what cell types they alter activity. Better understanding of how enhancer sequences drive activity patterns across cellular contexts would enable more accurate interpretation of the effects of non-coding mutations.

Enhancers harbor binding motifs recognized by TFs; thus the information encoded in enhancer sequences provides valuable information about regulatory specificity (2,9). Technological advances in high-throughput sequencing have enabled the development of genome-scale assays to identify sequences with putative enhancer activity. Several large-scale efforts have applied methods such as chromatin immunoprecipitation followed by sequencing (ChIP-seq) (10), identification of DNaseI-hypersensitive sites (DHS) via sequencing (DNase-seq) (11), and identification of enhancer RNA (eRNA) transcription via cap analysis of gene expression (CAGE) (12) to map putative enhancers over many tissues and cell lines (13–16).

Analyses of these and smaller-scale enhancer datasets have enabled identification of the unique sequence and chromatin properties of enhancers active in different tissues, which can then be used to predict enhancers in other contexts (17–19). Indeed, enhancer-finding algorithms based solely on sequence information have successfully predicted active enhancers in many tissues (20–24). These algorithms usually perform better than enhancer-finding algorithms built only on the occurrence profiles of known TF motifs, suggesting that the algorithms detect previously unidentified functional sequence characteristics that, if interpreted, could fill gaps in current knowledge about TF binding specificities and other enhancer sequence properties. For example, a recent study proposed a model in which short repetitive sequences—dinucleotide repeat motifs (DRMs)—promote general enhancer activity and play an essential role in driving broad enhancer activity across many cell types (16). In spite of these successes, we still lack a comprehensive understanding of how enhancer sequences drive their activity across tissues and development.

In this study, we comprehensively analyzed the of ability of short DNA sequence patterns, including DRMs, to predict the breadth of activity of tens of thousands of human enhancers across hundreds of human tissues. First, we computed the enrichment of DRMs among broadly active enhancers, and unlike in *Drosophila,* we consistently observed significant enrichment of GC DRMs and depletion of TA DRMs. To evaluate the ability of DRMs to predict broadly active enhancers, we trained a support vector machine (SVM) classification algorithm on the occurrence patterns of DRMs. In further contrast to results in *Drosophila,* we found that DRMs alone were only weakly predictive of broadly active enhancers versus context-specific enhancers or random regions from the genomic background. However, when trained on all possible 6-bp sequences, the algorithm could readily distinguish between broadly active, context-specific, and genomic background regions. The 6-mer sequence patterns most enriched—and therefore most predictive—of broadly active enhancers were GC-rich, suggesting that DRM contributions to enhancer activity are part of a larger trend seen among other 6-mers that is driven by GC content. Furthermore, we show that broadly active human TFs are more likely to bind GC-rich sequences than tissues-specific TFs. Thus, we conclude that, DRMs are not unique drivers of human enhancer activity, but broadly active human enhancers exhibit distinct sequence properties.

## METHODS

### Enhancer Data

We focused our analyses on enhancers identified by CAGE from the FANTOM Consortium across 411 different tissues and cellular contexts, which by definition exclude regions near known transcription start sites and exons of mRNAs (both protein-coding and noncoding) and lncRNAs (13). We subdivided their 38,538 robust enhancers based on the number of contexts in which an enhancer was found to be active. We defined the top 5% most active enhancers as the “broadly active” set; this corresponded to enhancers with activity in greater than 45 contexts.

We generated several sets of random non-enhancer regions for each enhancer set, using *shuffleBed* (25) to obtain length-matched regions for each input set of genomic regions. We also generated negative regions matched on GC content and chromosome as well as length using a custom script. We excluded locations in the positive set as well as all enhancers from the full permissive CAGE enhancer dataset (43,011 total sequences), ENCODE blacklist regions, genome (hg19) assembly gaps, and experimentally verified VISTA enhancers (downloaded in March 2014) (26) from the negatives. Further excluding regions near known transcription start sites and exons in addition did not materially change overall prediction performance (Figure S1), and there was a strong correlation between the weights assigned each 6-mer between the two classifiers (Spearman's ρ = 0.91, p ≈0), suggesting that they learned similar models of sequence.

To enable comparison with the fold enrichment analyses carried out by Yáñez-Cuna et al. (2014), we analyzed two additional human enhancer sets. We obtained DNase I hypersensitivity peaks and enhancer-associated histone modification data (15) from ENCODE (https://genome.ucsc.edu/ENCODE). Using *intersectBed* (25), we defined 13,069 broadly active DHS peaks found in at least 120 cell types, and 1,449 regions containing both H3K27ac and H3K4me1 marks that were active in at least 10 cell lines: GM12878, H1hesc, Hmec, Hsmm, Huvec, K562, Nha, Nhlf, Nhek, and Osteoblast. We also filtered both sets to exclude regions overlapping CpG islands from the CpG Islands track in the UCSC Genome Browser. Many of the DHS peaks are expected to be enhancers, but this set includes other regulatory regions as well. We generated matched negative regions for these sets using the criteria described above for CAGE enhancers.

### DRM Definition and Identification

We searched for DRMs using PWMs with probability of one for the appropriate nucleotide in each position: CACACA, GAGAGA, GCGCGC, and TATATA. We identified and counted DRM occurrences using the python package MOODS, which searches input DNA sequences on both strands for occurrences of motifs defined by PWMs (27,28), with a pseudocount of 0.001 and a match cutoff of *P* < 1/1024. Considering both strands meant that instances of the GC and TA repeats were counted twice, as they are their own reverse complements. We used the human genomic nucleotide frequencies for the background probabilities when calculating match scores and P-values, since the human genome is 42% GC.

We settled on these parameters after evaluating different combinations of thresholds and background frequencies with respect to the number and sequence diversity of DRMs we found. Using *P* < 1/4096 resulted in no perfect matches to the TA DRM passing the significance threshold, due to the higher genomic background frequency of TA bases (Figure S2). Thus, we chose *P <* 1/1024 as it minimized the number of inexact matches included, while still allowing all perfect matches to pass the cutoff. Results were similar when identifying only exact matches (Figure S3). We explicitly controlled for length in most analyses, because length is positively associated with activity in the FANTOM dataset. This step was unnecessary for relative fold enrichment analyses, as we compared relative occurrences in length-matched positive and negative sets.

Our parameters for defining DRMs differ from those used in the previous study, where they assumed an equal background probability for each nucleotide and used a PWM match cutoff of *P* < 1/256 (16,29). In addition, we used an invariant CA repeat motif, rather than the more variable motif inferred from STARR-seq data (Figure S2). We believe that considering the background human genome nucleotide frequencies is necessary, due to the non-uniform GC content genome-wide and in enhancers. We also chose to use a stricter threshold (P < 1/1024) for identifying matches to DRM motifs, because lower thresholds, such as 1/256, allowed many diverse, non-repetitive motifs to match. This is a partial cause of the lower DRM density we observed compared to Yáñez-Cuna et al. (2014). Additionally, using invariant motifs of consistent length and information content for all four DRMs facilitated direct comparison of the results for different DRMs. We felt that these settings best captured the notion of a “dinucleotide repeat motif.” Other than these differences, the parameters used were the same as in Yáñez-Cuna et al. as best as we could determine.

### Fold Enrichment Analyses

We calculated motif fold enrichment by dividing the mean count of the occurrence of the sequence in question for the enhancer set by that in the negative set, which was either the matched non-enhancer regions from the genomic background or the context-specific enhancers. When we were comparing enhancers to genomic backgrounds, we analyzed four independent negative sets separately, and then plotted the mean and standard deviation. P-values were calculated for the distribution of counts in broadly active enhancers vs. a negative set by the Wilcoxon rank sum test.

### Enhancer Prediction

To predict whether occurrence patterns of short DNA sequence motifs were sufficient to distinguish broadly active enhancers from the genomic background and from context-specific enhancers, we trained 6-mer spectrum kernel SVMs (30). The spectrum kernel is a string kernel that defines the similarity of two DNA sequences based on the occurrence of all possible short DNA sequence patterns of a given length, *k*, within them. We computed the weight given to each possible 6-mer by each SVM (31) and averaged the weights across training runs vs. four independent negative sets. For predictions using DRMs, we used the counts per base pair for each DRM as training features. To predict enhancers based on counts of motifs of known transcription factors, we used position weight matrices (PWMs) from the JASPAR 2016 vertebrate database (32) to count the occurrences of each motif in our genomic regions using FIMO under default settings (33). We then used the motif counts per base pair as features for the classifier.

Performance of all SVM classifiers was evaluated using 10-fold cross-validation, which limits overfitting by only training the classifier on a subset of the data at any given time. Receiver Operator Characteristic (ROC) and Precision Recall (PR) curves were calculated by averaging over the 10 cross-validation runs. All SVM analyses were performed using the SHOGUN Machine Learning Toolbox v4.0.0 (34). For the predictions of broadly active regions versus context-specific regions, we took a random subset of the larger set to maintain the number of regulatory regions considered across analyses. We controlled for length differences by expanding or contracting enhancers in each set to be 600 bp long while maintaining their original centers.

### Transcription factor binding motif and expression analysis

We obtained transcription factor binding motifs from the JASPAR 2016 vertebrate database (32). We obtained tissue specificity scores (TSPS) for 332 TFs from the FANTOM Consortium (35). A TF with uniform expression across all tissues is assigned a TSPS equal to zero, while a TF expressed in only a single tissue receives a maximum TSPS of ~5. Following the original analysis of TSPS, we classified 157 TFs as “specific” (TSPS ≥ 1) and 175 as “broad” that are expressed in a wider range of contexts (TSPS < 1). We compared the motif GC content distributions of the specific and broadly expressed TF groups using the Wilcoxon rank sum test.

## RESULTS

### DRMs are enriched (GC) and depleted (TA) in human enhancers, but the patterns do not match those in *Drosophila*

Recent work in *Drosophila* suggested that DRMs are a general feature of enhancers and that presence of many DRMs in an enhancer is a main driver of broad regulatory activity across diverse tissues (16). To test the hypothesis that high DRM occurrence drives broad enhancer activity across tissues in humans (16), we analyzed sequence patterns in putative enhancers across diverse human cells and tissues. We considered 38,538 transcribed enhancers identified via CAGE for 411 contexts by the FANTOM consortium (13). We defined the 1961 enhancers in the top 5% of the breadth of activity distribution (active in more than 46 contexts) as broadly active.

As a first step in investigating the contribution of DRMs to human enhancer activity, we computed the relative enrichment of DRMs in broadly active enhancers compared to context-specific enhancers and length-matched background regions using position weight matrices (PWMs). *Drosophila* enhancers exhibit enrichment for all DRMs except TA, and also show a positive association between DRM frequency and breadth of activity most strongly with GA and CA repeats, and GC to a lesser extent. Thus, under the *Drosophila* DRM model, we would expect CA, GA, and GC DRMs to be enriched in broadly active human enhancers compared to the other sets.

In humans, the CA, GA, and GC DRMs were all significantly enriched in broadly active enhancers compared to the genomic background (Figure 1A; *P* = 4.1E–16, *P* = 2.0E–14, and *P* < 2.2E–16, respectively). However, the magnitude of the enrichments for CA (1.2x) and GA (1.7x) were modest compared to GC (11.8x), and when compared to context-specific enhancers, significant enrichment remained only for the GC DRM (Figure 1B). The TA DRM, on the other hand, was significantly depleted compared to both the genomic background (–3.9x, *P* = 3.0E–7) and context-specific enhancers (–2.9x, *P* = 2.2E–16).

**Figure 1.**
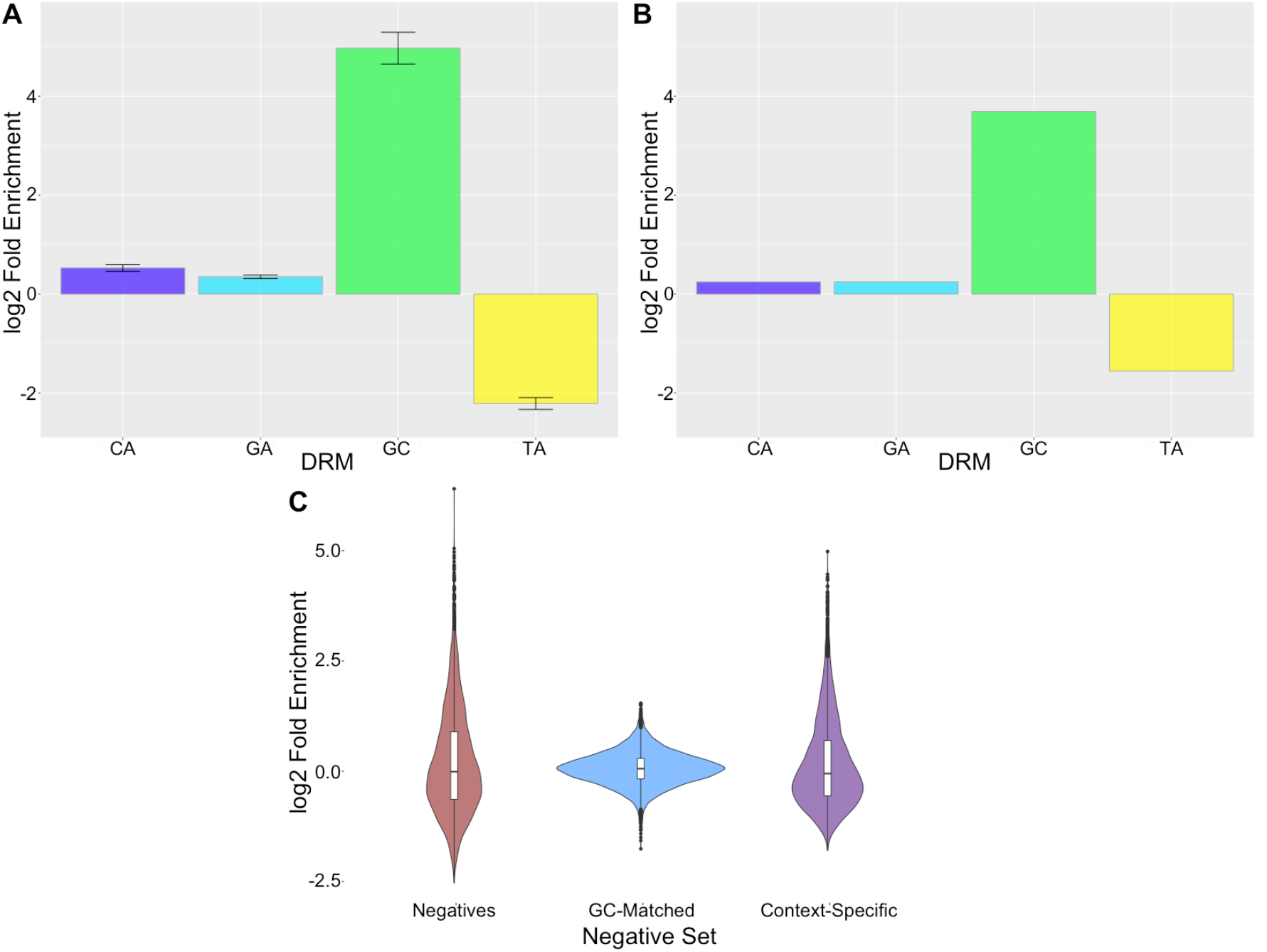
GC DRMs are enriched and TA DRMs depleted in broadly active enhancers. log_2_(Fold Enrichment) of the occurrence of each DRM in broadly active enhancers vs. (A) genomic background and (B) context-specific enhancers. Error bars are standard errors over four replicates. (C) log_2_(Fold Enrichment) of all 6-mers in broadly active enhancers vs. genomic background (red), GC-matched background (blue) or context-specific enhancers (purple).

Furthermore, GC DRM density (DRM/bp) significantly increased as breadth of activity increased (Figure S4; Spearman's ρ = 0.12, *P* ≈ 0), indicating that this is a general trend across enhancers. Similarly, TA DRM density significantly decreased with breadth of activity (Spearman's ρ = −0.020; *P* = 9.72E-05). There was not a significant trend for GA and CA DRMs (Figure S4).

To confirm that the observed trends in DRM patterns were not unique to the transcribed enhancers defined by CAGE, we also analyzed DRM patterns in a “histone-derived set” of 1,449 enhancers, consisting of regions with overlapping H3K4me1 and H3K27ac histone marks from 13 contexts (36), and a “DHS set,” of 13,069 DNaseI hypersensitive peaks across 126 contexts from ENCODE (15). The DRM enrichment patterns were similar in these enhancer sets to those observed for CAGE enhancers: GC was significantly enriched and TA significantly depleted (Figure S5).

The majority of the broadly active histone-mark-defined enhancers contained at least one DRM (Figure S5); this is in contrast to their relative rarity in the CAGE and DHS sets. The increased counts are likely due to the greater length (and presumably lower resolution) of the histone-derived set: average enhancer length of 5,797 bp vs. 200 and 297 bp for the DHS and CAGE sets, respectively. In contrast, the *Drosophila* enhancers were 500 bp long and had median DRM counts between 1 and 6, which is more similar to the histone set despite being an order of magnitude smaller (16,37).

Overall, DRM patterns in human enhancers do not match the patterns observed for *Drosophila* enhancers. However, it is possible that DRMs in general maintain importance in driving broad enhancer activity between these diverse species, but the specific motifs are not conserved.

### DRMs alone are weakly predictive of broadly active human enhancers

To directly evaluate the ability of DRMs to identify broadly active human enhancers, we used a support vector machine (SVM) learning framework (17). We trained a linear SVM classifier to distinguish broadly active enhancers from context-specific enhancers and the genomic background using patterns of DRM occurrence. Using only DRM counts as features yielded poor performance at each classification task (Figures 2A and S6A). We first trained the SVM to distinguish broadly active enhancers from a set of length-matched genomic background regions that excluded all putative enhancers, gaps in the genome assembly, and ENCODE blacklist regions (Methods). In 10-fold cross validation, the classifier achieved an area under the receiver operating characteristic curve (ROC AUC) of 0.61 and a precision recall (PR) AUC of 0.64. Enhancers are known to have greater GC content compared to the genomic background (mean 46% vs. 42%), so to determine whether DRM sequence patterns held predictive value independent from overall GC content, we repeated the previous analysis training the classifier on negative training sets generated from random background regions matched to broadly active enhancers for both length and GC content. This classifier had a drop in PR AUC compared to the non-GC-matched classifier (ROC AUC = 0.55, PR AUC = 0.54), suggesting that GC content was important for some for the predictive ability of DRMs.

Next, we evaluated the ability of DRMs to distinguish broadly active enhancers from 1961 context-specific enhancers (Figure 2A). Since the context-specific enhancers were shorter on average (Figure S7), we controlled for length by expanding or contracting all enhancers in both sets to be 600 bp long, approximately the mean length of the most active enhancers. This DRM-based classifier trained vs. context-specific enhancers had a ROC AUC of 0.56 and PR AUC of 0.61. Because DRMs were rare in broadly active enhancers (median occurrence of zero for all DRMs; Figure 2B), the poor performance of the DRM-based SVM is not surprising. This suggests that DRMs are not major drivers of enhancer activity in humans.

**Figure 2.**
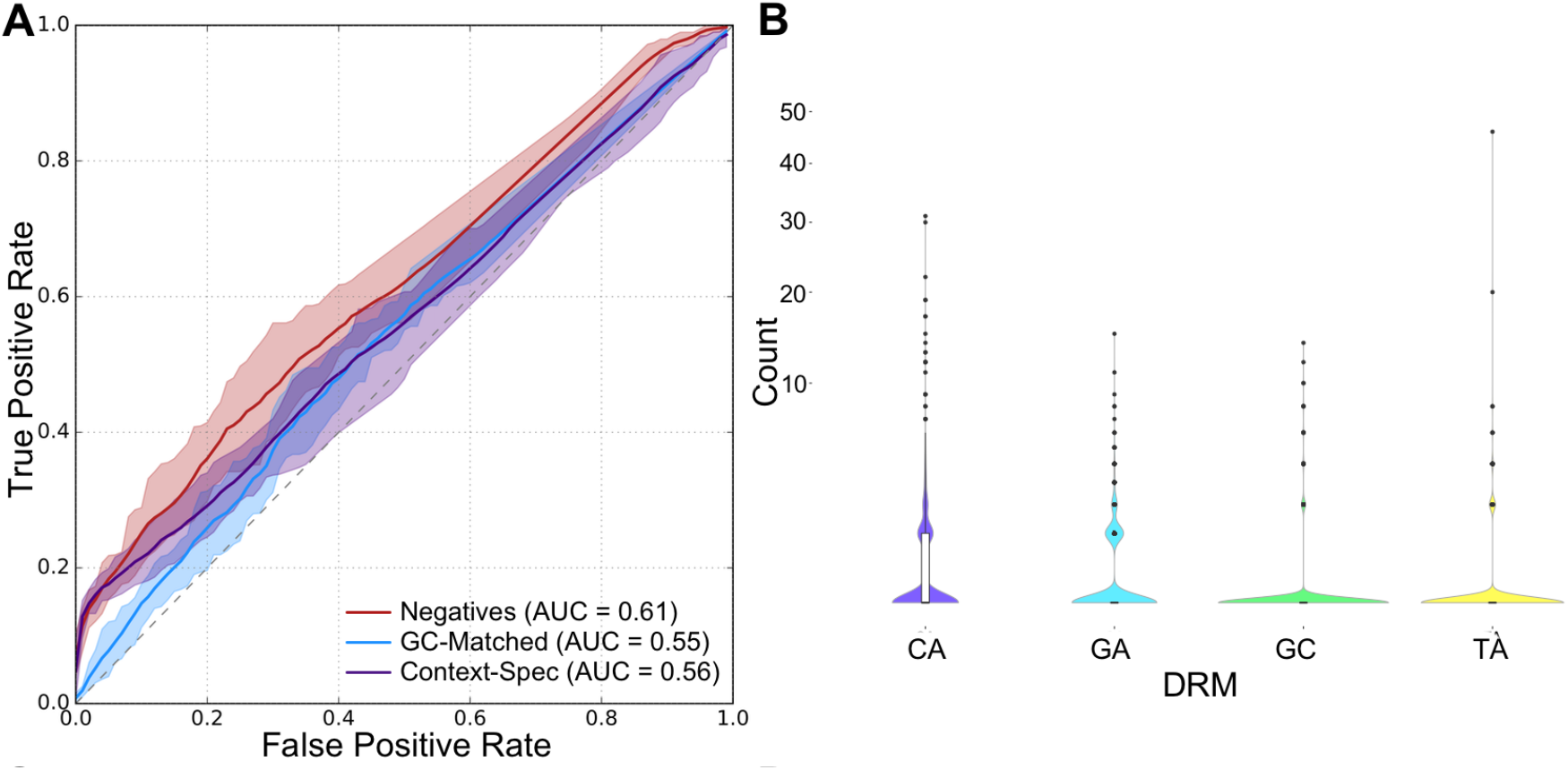
DRMs are not major drivers of human broadly active human enhancers. (A) ROC curves for SVM-based classification of broadly active enhancers vs. length-matched genomic background (red), GC-matched genomic background (blue), and context-specific enhancers (purple) using the frequency of the four DRMs as features. (B) The distribution of occurrences for each DRM observed over all broadly active enhancers. Most enhancers do not contain each class of DRM. The area under each curve (AUC) is given in parentheses. Shaded areas are bounded by the maximum and minimum observed ROC. Precision-recall curves are given in Figure S6. Box plots show median and 1^st^/3^rd^ quartiles.

### Comprehensive analysis of short DNA sequence motif occurrence accurately identifies broadly active human enhancers

Given that DRMs by themselves were only weakly predictive of broadly active human enhancers, we evaluated the ability of additional short DNA sequence motifs to predict the breadth of enhancer activity. Using the occurrence patterns of all 4,096 possible 6-mers in the enhancer sequence as features in a spectrum kernel SVM (30), we repeated the classifications performed for the DRMs. The classifier trained on broadly active enhancers vs. random background regions performed very well (Figure 3A; ROC AUC = 0.93, PR AUC = 0.92). When classifying GC-matched regions, the performance of this classifier decreased, but was still strong (Figure 3A; ROC AUC = 0.87, PR AUC = 0.86). Furthermore, the classifier performed as well at distinguishing between broadly active and context-specific enhancer classes as it did distinguishing broadly active enhancers from genomic background (Figures 3A and S6B; ROC AUC = 0.87, PR AUC = 0.88).

**Figure 3.**
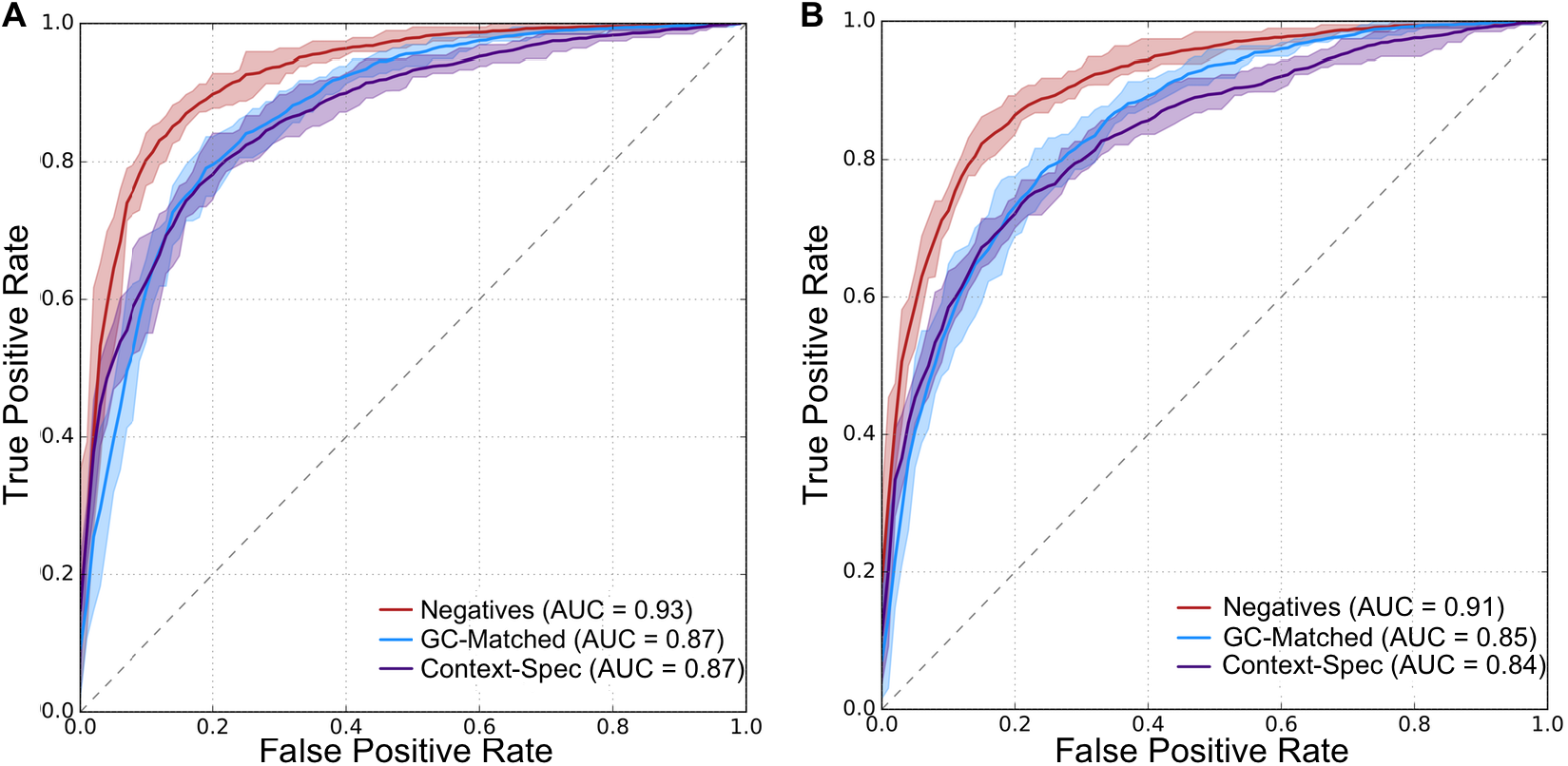
Short DNA sequence patterns accurately distinguish broadly active human enhancers from the genomic background and context-specific enhancers. (A) Classifiers trained using all possible 6-mers or (B) density of TF motifs as features. ROC curves were calculated using 10-fold cross-validation and averaging the ROC obtained by each round of validation. The area under each curve (AUC) is given in parentheses. Shaded areas are bounded by the maximum and minimum observed ROC. Precision-recall curves are given in Figure S6.

To test whether considering all possible 6-mers increased performance compared to using current knowledge of TF binding preferences, we evaluated the performance of classifiers trained using counts of matches to 332 known TF binding motifs. All three classifiers performed slightly worse in these analyses than when considering all 6-mers (Figure 3A vs. 3B); however, the difference was most pronounced when distinguishing broadly active enhancers from context-specific enhancers (Figures 3B and S6C; ROC AUC = 0.84, PR AUC = 0.85).Thus, limiting the training features to current knowledge of TF specificity modestly decreased performance.

Since the relative performance of the classifiers indicates that DRMs only contribute modestly to enhancer sequence activity patterns, we evaluated their contribution to human enhancer activity in the context of all possible 6-mers (Figure 1C). The enrichment of the GC DRM in broadly active enhancers was more than two standard deviations (SDs) above the mean over all 6-mer enrichments for all three comparisons. The TA DRM was more than 1 SD less than the mean for the broadly active enhancers vs. genomic background and context-specific enhancers (Figure 1C). The CA and GA DRMs were both within 1 SD of the mean for all three comparisons.This suggests that the GC DRM, and to a lesser extent the TA DRM are enriched and depleted, respectively, in broadly enhancers compared to the enrichment of 6-mers in general. Despite this enrichment, the rarity of DRMs in broadly active enhancers (Figure 2B) reduced their predictive ability overall. Collectively, these results show that DRMs alone are not nearly as informative about enhancer activity and breadth as models that include additional short sequence patterns or known TF binding motifs.

### GC-rich motifs are predictive of broadly active enhancers

Given the elevated GC content of enhancers and the enrichment and depletion of the GC and TA DRMs (the two DRMs with unequal GC content), we quantified the relationship between GC content and 6-mer enrichment in broadly active enhancers. In comparisons with the genomic background, the correlation was significantly positive (Figure 4A; Spearman's ρ = 0.87, *P* ≈ 0). This is not surprising given that enhancers have high GC content compared to the genomic background. As expected, this trend was strongly attenuated in the GC-matched comparison (Figure 4B; Spearman's ρ = 0.045, *P* = 0.004).

**Figure 4.**
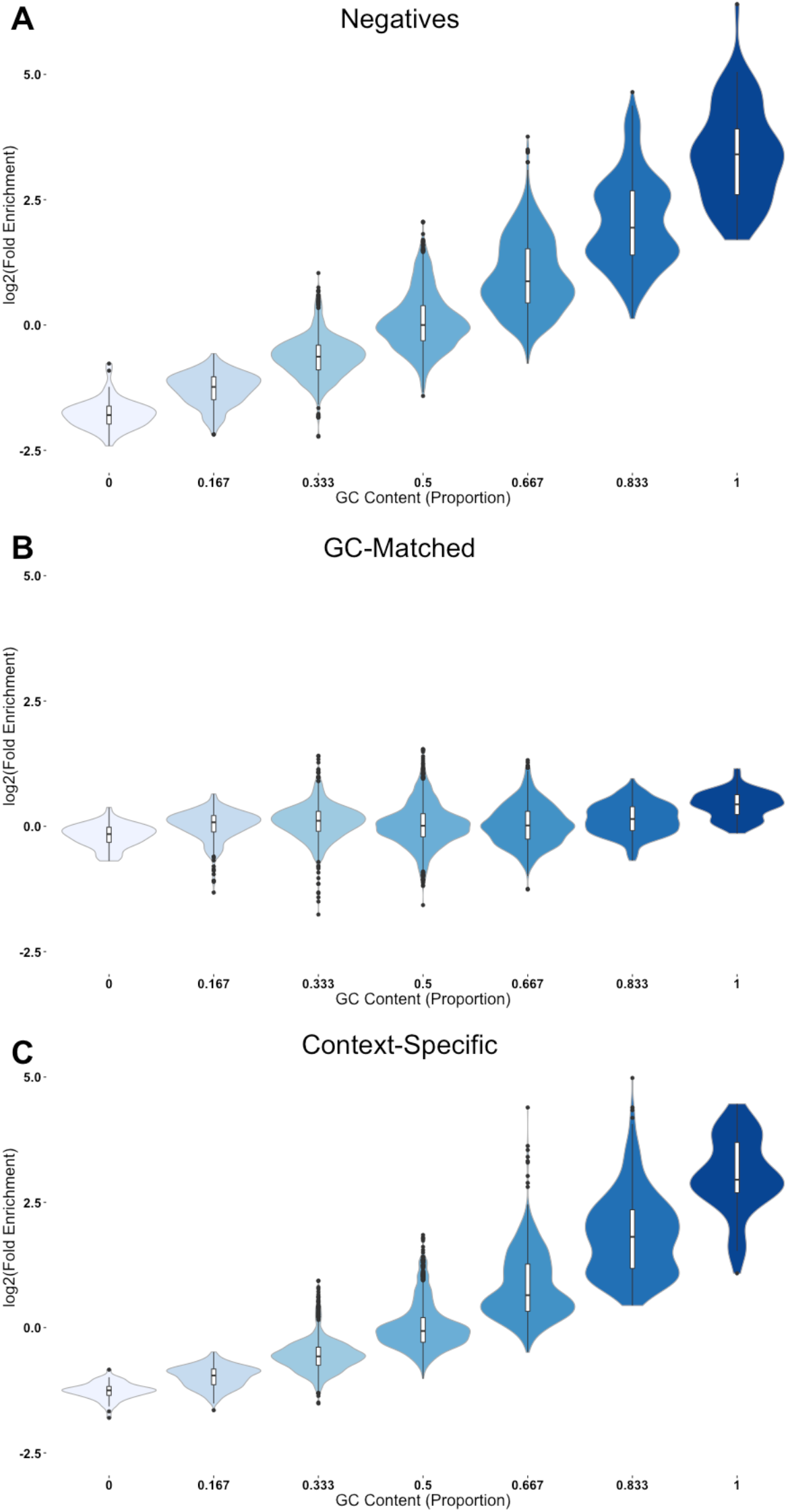
Short DNA sequence patterns enriched in broadly active enhancers have high GC content. The log_2_ of the fold enrichment for each 6-mer is plotted against the GC-content of the 6-mer for comparisons of broadly active vs. (A) non-GC-matched background, (B) the GC-matched background, and (C) context-specific enhancers. Means were taken over the enrichments compared to four different background sets.

We previously observed that enhancer GC content varied in different tissues’ enhancers (17), and here we found that GC content is positively correlated with breadth of activity among the enhancers (Figure S8; Spearman's ρ = 0.25, *P* ≈ 0). Similarly, GC content showed a high correlation with enrichment in broadly active vs. context-specific enhancers (Figure 4C; Spearman's ρ = 0.88, *P* ≈ 0). This mirrors the patterns shown by the GC and TA DRMs.

The classification function learned by a trained spectrum kernel SVM implicitly assigns weights to each 6-mer that indicate its contribution to the classifier's prediction. Repeating the GC content analyses using these 6-mer weights rather than their enrichment resulted in similar correlations (Figure S9; Spearman's ρ = 0.31, 0.014, 0.29, *P* ≈ 0 for genomic background, GC-matched, and context-specific enhancers respectively). This argues that, in terms of both individual motif enrichment and importance to trained classifiers, high GC content is characteristic of broadly active enhancers, regardless of status as a DRM.

### Broadly active TFs have GC-rich motifs

The highly weighted/enriched motifs likely serve important biological functions that contribute to enhancer activity. Since enhancers function by binding transcription factors, we hypothesized that DNA sequence patterns that facilitate the binding of broadly expressed transcription factors could drive broad enhancer activity across many contexts. To explore this, we analyzed the sequences and breadth of expression of known TF binding motifs from the JASPAR database (32). We classified the TFs into broadly expressed and tissue-specific classes based on expression data from the FANTOM Consortium (35). In support of our hypothesis, the motifs of broadly active TFs have significantly higher GC content than those of context-specific TFs (Figure 5; *P* = 2.4E-8), mirroring the trend seen in 6-mers predictive of broad enhancer activity.

**Figure 5.**
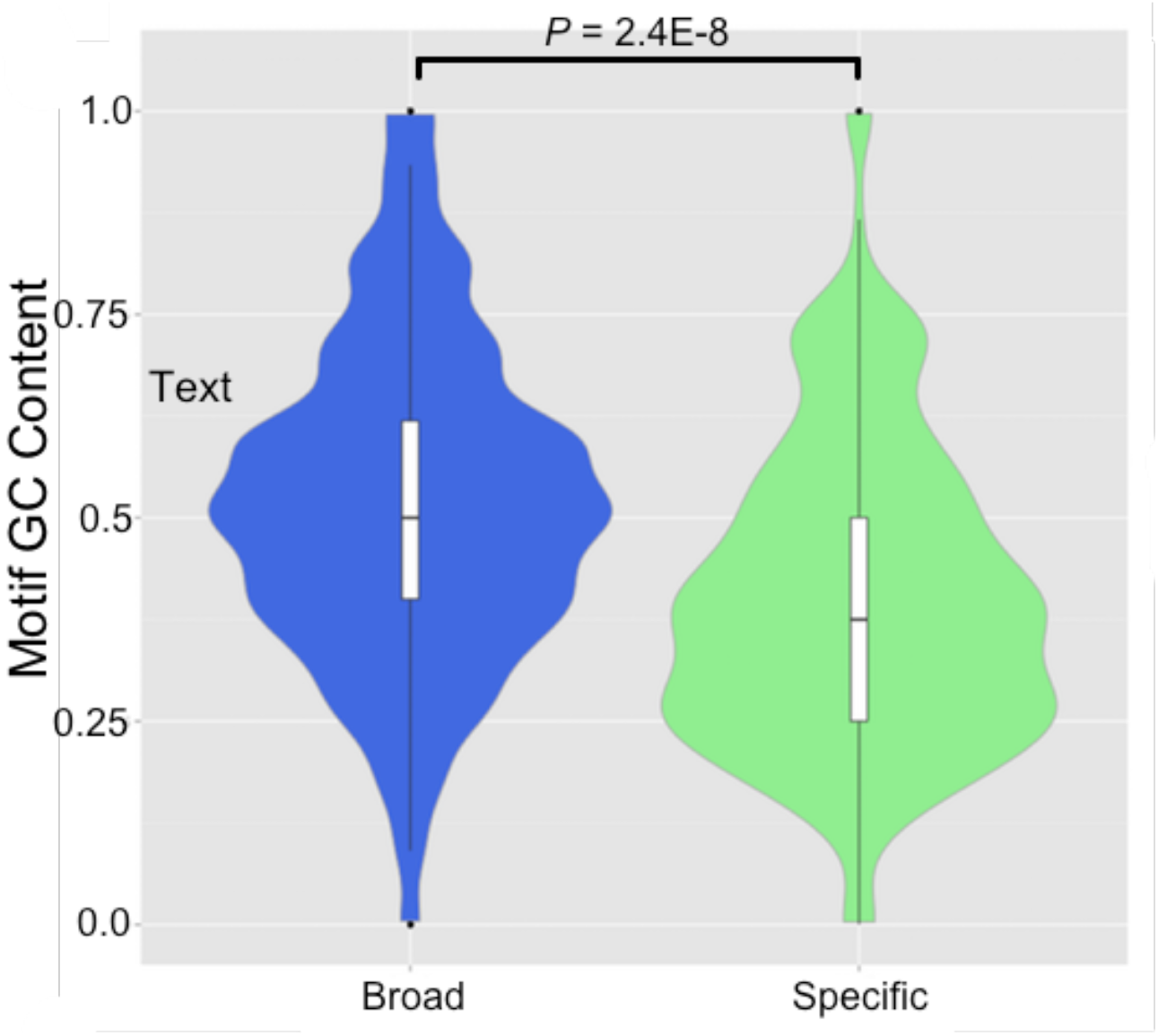
Binding motifs for broadly expressed TFs are more GC-rich than motifs for context-specific TFs. The median GC content (0.50) of motifs recognized by 175 broadly expressed TFs (blue) was significantly (*P* = 2.4E-8) greater than the motif GC content (0.38) of context-specific TFs (green). Box plots show median, and 1^st^/3^rd^ quartile.

## DISCUSSION

We analyzed the contribution of DRMs and other short DNA sequence motifs to the activity patterns of human enhancers across hundreds of cellular contexts. In contrast to the model proposed in *Drosophila* (16), GC DRMs were enriched in broadly active enhancers, while TA DRMs were depleted. Using an unbiased machine learning framework, we found that DRM occurrence patterns were only weakly predictive of broadly active human enhancers (ROC AUC ranging from 0.55 to 0.61). However, a classifier trained on the occurrence of all possible 6-bp sequences very accurately distinguished broadly active human enhancers from the genomic background (ROC AUC = 0.93), GC-matched background regions (ROC AUC = 0.87), and context-specific enhancers (ROC AUC = 0.87). Furthermore, 6-mers highly predictive of broad activity tended to be GC-rich, while those with the most negative weights tended to be GC-poor, even when classifying GC-matched regions. These results suggest that broadly active enhancers have distinct sequence properties, and that the enrichment and depletion of DRM sequences is part of a larger pattern in which particularly GC-rich and GC-poor sequences are indicative of broad and context-specific activity, respectively. Consistent with this pattern, TFs with broad expression have greater affinity for GC-rich motifs than TFs with tissue-specific expression patterns.

Our findings in human enhancers differ from recent results in *Drosophila* in several respects. Broadly active *Drosophila* enhancers exhibit enrichment for all DRMs except TA, while broadly active human enhancers are consistently enriched only for the GC DRM. We also found that DRM counts alone are significantly less predictive of enhancer activity than wider sequence patterns or *Drosophila* models including many motifs (16). Other sequences predictive of broad enhancer activity tend to be GC-rich, demonstrating that the effects on human enhancer sequence activity are not unique to repeat elements.

There are several possible causes of the observed differences in DRM patterns between humans and *Drosophila*. First, they could be due to differences in the enhancer identification strategy used. The main set of human enhancers analyzed was identified using CAGE to detect native eRNAs, while the *Drosophila* enhancer sets were assembled using STARR-seq (37). Both methods have potential weaknesses. CAGE-seq is only able to identify enhancers that produce bidirectional capped transcripts, while the STARR-seq assay isolates potential regulatory sequences in reporter constructs separate from their genomic contexts and thus could introduce activity patterns not representative of enhancers in their natural chromatin context. To address this concern, we analyzed other human sets defined using functional genomics data (histone modifications and DNaseI hypersensitivity data). We found patterns consistent with the CAGE enhancers, so this suggests that our findings are robust among human enhancers. Second, differences in the number of biological contexts considered could influence the comparison. We considered enhancer activity across 411 human cellular contexts, while only three cell types were considered in the *Drosophila* study. These cells were from different lineages and developmental stages, but further work that considers more cellular contexts in *Drosophila* would be necessary for a more direct comparison. Finally, there were a number of technical differences in how DRMs were defined between the studies. For example, we used stricter P-value thresholds for calling DRMs and a background model tailored to the genome GC content rather than uniform frequencies. We felt that these definitions better reflected the concept of a “dinucleotide repeat motif’ and enabled comparison between different motifs. Nonetheless, we found that this and other technical differences did not dramatically influence our results (Figures S1-3).

Thus, while technical factors may have contributed, the observed differences were likely also influenced by biological differences between the *Drosophila* and human genomes. For example, despite having similar GC content, the *Drosophila* genome is not as CpG-depleted as humans (38). This could influence the roles and dynamics of CpG islands in enhancer activity between the species. In addition, while recent studies of transcriptional networks and TF binding preferences have revealed remarkable conservation of elements of metazoan gene regulation (39–41), there are considerable differences in the TF complement and gene expression patterns between these two species. It is possible that the differences in DRM enrichments reflect a difference in the sets of TFs that bind broadly active enhancers in the two species, or that broadly expressed transcription factors in *Drosophila* do not show the same collectively higher GC content compared to context-specific TFs (Figure 5). The differences in the role of DRMs and other regulatory sequence motifs between humans, flies, and other animals must be explored further, but such studies will require comprehensive catalogs of enhancers active across many tissues in additional species.

In conclusion, we demonstrate that while short DNA sequence patterns can accurately identify broadly active human enhancers, DRMs are not the main drivers of activity. This emphasizes the importance of DNA sequence patterns on enhancer biology beyond existing knowledge of transcription factor binding motifs, and suggests several avenues for future research. Most importantly, more work is needed to understand the regulatory logic of enhancer sequences; we suspect that highly predictive sequence patterns could be mined to identify novel binding motifs and combinatorial interactions. Our results also reveal that we understand relatively little about how enhancer sequence and activity evolve. Resolving the evolution and mechanistic functions of these enriched sequences will require further statistical and experimental analyses, but the approach presented here provides a framework in which to quantify and explore how DNA sequence influences gene regulatory activity.

## ACKNOWLEDGEMENTS

We thank Alexandra Fish, Emily Hodges, and Corinne Simonti for helpful discussions and comments on the manuscript.

## FUNDING

L.L.C. was supported by the National Institutes of Health [T32GM080178]. J.A.C. was supported by the National Institutes of Health [1R01GM115836], a March of Dimes Innovation Catalyst award, and institutional funds from Vanderbilt University.

## REFERENCES

1. Levine, M. (2010) Transcriptional Enhancers in Animal Development and Evolution. Current Biology, 20, R754–R763.

2. Shlyueva, D., Stampfel, G. and Stark, A. (2014) Transcriptional enhancers: from properties to genome-wide predictions. Nat Rev Genet, 15, 272–286.

3. Corradin, O. and Scacheri, P.C. (2014) Enhancer variants: evaluating functions in common disease. Genome Medicine, 6, 85.

4. Maurano, M.T., Humbert, R., Rynes, E., Thurman, R.E., Haugen, E., Wang, H., Reynolds, A.P., Sandstrom, R., Qu, H., Brody, J. et al. (2012) Systematic Localization of Common Disease-Associated Variation in Regulatory DNA. Science, 337, 1190–1195.

5. Bauer, D.E., Kamran, S.C., Lessard, S., Xu, J., Fujiwara, Y., Lin, C., Shao, Z., Canver, M.C., Smith, E.C., Pinello, L. et al. (2013) An Erythroid Enhancer of BCL11A Subject to Genetic Variation Determines Fetal Hemoglobin Level. Science, 342, 253–257.

6. Fortini, B.K., Tring, S., Plummer, S.J., Edlund, C.K., Moreno, V., Bresalier, R.S., Barry, E.L., Church, T.R., Figueiredo, J.C. and Casey, G. (2014) Multiple Functional Risk Variants in a SMAD7 Enhancer Implicate a Colorectal Cancer Risk Haplotype. PLoS ONE, 9, e111914.

7. Smemo, S., Tena, J.J., Kim, K.-H., Gamazon, E.R., Sakabe, N.J., Gomez-Marin, C., Aneas, I., Credidio, F.L., Sobreira, D.R., Wasserman, N.F. et al. (2014) Obesity-associated variants within FTO form long-range functional connections with IRX3. Nature, 507 371–375.

8. Musunuru, K., Strong, A., Frank-Kamenetsky, M., Lee, N.E., Ahfeldt, T., Sachs, K.V., Li, X., Li, H., Kuperwasser, N., Ruda, V.M. et al. (2010) From noncoding variant to phenotype via SORT1 at the 1p13 cholesterol locus. Nature, 466, 714–719.

9. Cheng, C., Alexander, R., Min, R., Leng, J., Yip, K.Y., Rozowsky, J., Yan, K.-K., Dong, X., Djebali, S., Ruan, Y. et al. (2012) Understanding transcriptional regulation by integrative analysis of transcription factor binding data. Genome Research, 22, 1658–1667.

10. Johnson, D.S., Mortazavi, A., Myers, R.M. and Wold, B. (2007) Genome-Wide Mapping of in Vivo Protein-DNA Interactions. Science, 316, 1497–1502.

11. Boyle, A.P., Davis, S., Shulha, H.P., Meltzer, P., Margulies, E.H., Weng, Z., Furey, T.S. and Crawford, G.E. (2008) High-Resolution Mapping and Characterization of Open Chromatin across the Genome. Cell, 132, 311–322.

12. Shiraki, T., Kondo, S., Katayama, S., Waki, K., Kasukawa, T., Kawaji, H., Kodzius, R., Watahiki, A., Nakamura, M., Arakawa, T. et al. (2003) Cap analysis gene expression for high-throughput analysis of transcriptional starting point and identification of promoter usage. Proceedings of the National Academy of Sciences, 100, 15776–15781.

13. Andersson, R., Gebhard, C., Miguel-Escalada, I., Hoof, I., Bornholdt, J., Boyd, M., Chen, Y., Zhao, X., Schmidl, C., Suzuki, T. et al. (2014) An atlas of active enhancers across human cell types and tissues. Nature, 507, 455–461.

14. Kundaje, A., Meuleman, W., Ernst, J., Bilenky, M., Yen, A., Heravi-Moussavi, A., Kheradpour, P., Zhang, Z., Wang, J., Ziller, M.J. et al. (2015) Integrative analysis of 111 reference human epigenomes. Nature, 518, 317–330.

15. Thurman, R.E., Rynes, E., Humbert, R., Vierstra, J., Maurano, M.T., Haugen, E., Sheffield, N.C., Stergachis, A.B., Wang, H., Vernot, B. et al. (2012) The accessible chromatin landscape of the human genome. Nature, 489, 75–82.

16. Yáñez-Cuna, J.O., Arnold, C.D., Stampfel, G., Boryn, L.M., Gerlach, D., Rath, M. and Stark, A. (2014) Dissection of thousands of cell type-specific enhancers identifies dinucleotide repeat motifs as general enhancer features. Genome Research, 24, 1147–1156.

17. Erwin, G.D., Oksenberg, N., Truty, R.M., Kostka, D., Murphy, K.K., Ahituv, N., Pollard, K.S. and A., C.J. (2014) Integrating diverse datasets improves developmental enhancer prediction. PLOS Computational Biology, 10.

18. Capra, J.A. (2015) Extrapolating histone marks across developmental stages, tissues, and species: an enhancer prediction case study. BMC Genomics, 16, 1–9.

19. Ernst, J. and Kellis, M. (2015) Large-scale imputation of epigenomic datasets for systematic annotation of diverse human tissues. Nat Biotech, 33, 364–376.

20. Lee, D., Karchin, R. and Beer, M.A. (2011) Discriminative prediction of mammalian enhancers from DNA sequence. Genome Res, 21, 2167–2180.

21. Burzynski, G.M., Reed, X., Taher, L., Stine, Z.E., Matsui, T., Ovcharenko, I. and McCallion, A.S. (2012) Systematic elucidation and in vivo validation of sequences enriched in hindbrain transcriptional control. Genome Res, 22, 2278–2289.

22. Ghandi, M., Lee, D., Mohammad-Noori, M. and Beer, M.A. (2014) Enhanced Regulatory Sequence Prediction Using Gapped *k*-mer Features. PLoS Comput Biol, 10, e1003711.

23. Taher, L., Narlikar, L. and Ovcharenko, I. (2012) CLARE: Cracking the LAnguage of Regulatory Elements. Bioinformatics, 28, 581–583.

24. Narlikar, L., Sakabe, N.J., Blanski, A.A., Arimura, F.E., Westlund, J.M., Nobrega, M.A. and Ovcharenko, I. (2010) Genome-wide discovery of human heart enhancers. Genome Research, 20, 381–392.

25. Quinlan, A.R. and Hall, I.M. (2010) BEDTools: a flexible suite of utilities for comparing genomic features. Bioinformatics, 26, 841–842.

26. Visel, A., Minovitsky, S., Dubchak, I. and Pennacchio, L.A. (2007) VISTA Enhancer Browser—a database of tissue-specific human enhancers. Nucleic Acids Research, 35, D88–D92.

27. Korhonen, J., Martinmäki, P., Pizzi, C., Rastas, P. and Ukkonen, E. (2009) MOODS: fast search for position weight matrix matches in DNA sequences. Bioinformatics, 25, 3181–3182.

28. Pizzi, C., Rastas, P. and Ukkonen, E. (2011) Finding Significant Matches of Position Weight Matrices in Linear Time. Computational Biology and Bioinformatics, IEEE/ACM Transactions on, 8, 69–79.

29. Yáñez-Cuna, J.O., Dinh, H.Q., Kvon, E.Z., Shlyueva, D. and Stark, A. (2012) Uncovering cis-regulatory sequence requirements for context-specific transcription factor binding. Genome Research, 22, 2018–2030.

30. Leslie, C., Eskin, E. and Noble, W.S. (2002) The spectrum kernel: a string kernel for SVM protein classification. Pac Symp Biocomput, 564–575.

31. Guyon, I., Weston, J., Barnhill, S. and Vapnik, V. (2002) Gene Selection for Cancer Classification using Support Vector Machines. Mach. Learn., 46, 389–422.

32. Mathelier, A., Zhao, X., Zhang, A.W., Parcy, F., Worsley-Hunt, R., Arenillas, D.J., Buchman, S., Chen, C.-y., Chou, A., Ienasescu, H. et al. (2013) JASPAR 2014: an extensively expanded and updated open-access database of transcription factor binding profiles. Nucleic Acids Research.

33. Grant, C.E., Bailey, T.L. and Noble, W.S. (2011) FIMO: scanning for occurrences of a given motif. Bioinformatics, 27, 1017–1018.

34. Sonnenburg, S., Ratsch, G., Henschel, S., Widmer, C., Behr, J., Zien, A., Bona, F.d., Binder, A., Gehl, C. and Franc, V. (2010) The SHOGUN Machine Learning Toolbox. J. Mach. Learn. Res., 11, 1799–1802.

35. Ravasi, T., Suzuki, H., Cannistraci, C.V., Katayama, S., Bajic, V.B., Tan, K., Akalin, A., Schmeier, S., Kanamori-Katayama, M., Bertin, N. et al. (2010) An atlas of combinatorial transcriptional regulation in mouse and man. Cell, 140, 744–752.

36. ENCODE Project Consortium. (2012) An integrated encyclopedia of DNA elements in the human genome. Nature, 489, 57–74.

37. Arnold, C.D., Gerlach, D., Stelzer, C., Boryn, L.M., Rath, M. and Stark, A. (2013) Genome-Wide Quantitative Enhancer Activity Maps Identified by STARR-seq. Science, 339, 1074–1077.

38. Jabbari, K. and Bernardi, G. (2004) Cytosine methylation and CpG, TpG (CpA) and TpA frequencies. Gene, 333, 143–149.

39. Boyle, A.P., Araya, C.L., Brdlik, C., Cayting, P., Cheng, C., Cheng, Y., Gardner, K., Hillier, L.W., Janette, J., Jiang, L. et al. (2014) Comparative analysis of regulatory information and circuits across distant species. Nature, 512, 453–456.

40. Nitta, K.R., Jolma, A., Yin, Y., Morgunova, E., Kivioja, T., Akhtar, J., Hens, K., Toivonen, J., Deplancke, B., Furlong, E.E.M. et al. (2015) Conservation of transcription factor binding specificities across 600 million years of bilateria evolution. eLife, 4, e04837.

41. Gerstein, M.B., Rozowsky, J., Yan, K.-K., Wang, D., Cheng, C., Brown, J.B., Davis, C.A., Hillier, L., Sisu, C., Li, J.J. et al. (2014) Comparative analysis of the transcriptome across distant species. Nature, 512, 445–448.

